# Discovering conserved regulatory modules in predicted gene regulatory networks across species

**DOI:** 10.64898/2026.05.15.725337

**Authors:** Jingyi Zhang, Lenwood S. Heath

## Abstract

The discovery of conserved regulatory motifs across different species is a fundamental challenge in systems biology, especially considering the noisy and incomplete nature of predicted gene regulatory networks (GRNs) and the intractability of the underlying graph alignment problem. Traditional network alignment methods frequently enforce one-to-one node mappings or strict topological isomorphism, which fail to accommodate the many-to-many orthology mappings caused by evolutionary gene duplication. Consequently, strict constraints often yield highly fragmented topological islands rather than cohesive functional modules. In this work, we propose a relaxed topological alignment algorithm designed to extract conserved regulatory structures from cross-species GRNs. We formulate the discovery process as a multi-objective optimization problem that balances sequence homology, functional coherence, and a normalized topological consensus. To navigate the exponentially scaling search space, we introduce a greedy seed-and-extend heuristic bounded by a dynamic *ϵ*-stopping condition, which evaluates marginal objective gains to prevent functional dilution. We validate our algorithm using time-series transcriptomic data from *Arabidopsis thaliana, Zea mays*, and *Sorghum bicolor* focused on drought and developmental stress responses. While a strict topological baseline extracted only fragmented subgraphs limited to 51 homologous tuples, our relaxed heuristic successfully converged on a highly connected 444-tuple module. The resulting topology effectively links strictly conserved upstream transcription factors to their highly duplicated, species-specific downstream pathways. Our algorithm provides a robust, scalable computational methodology for identifying core regulatory logic across complex biological systems, facilitating the translation of conserved network architectures among multiple species.

**Author summary:** Identifying shared regulatory mechanisms across diverse species is essential for understanding how complex biological systems evolve and adapt. However, traditional computer algorithms struggle to align these biological networks because evolution frequently duplicates genes, breaking simple one-to-one comparisons and producing highly fragmented results. To overcome this limitation, we developed a relaxed cross-species network alignment algorithm. Instead of demanding perfectly identical network shapes, our approach dynamically balances genetic sequence similarity, network structure, and biological function. We demonstrated the performance of our algorithm using plant drought-stress networks as a case study. While strict methods only found tiny, disconnected network fragments, our algorithm uncovered a functionally coherent, interconnected regulatory module across three distinct species. We discovered that while upstream command genes remain strictly conserved, they regulate highly customized, species-specific execution pathways downstream. Ultimately, our framework provides a scalable, species-agnostic method to decode complex systems, allowing researchers to translate conserved biological logic across diverse genomes.

## Introduction

The inference and analysis of gene regulatory networks (GRNs) provide a powerful computational algorithm for understanding how collective systemic functions emerge from individual molecular components [1, 2]. As high-throughput transcriptomic data scales, graph-based abstractions have become essential for modeling these complex systems, where nodes represent genes and edges denote regulatory dependencies. A fundamental objective in comparative systems biology is the identification of conserved regulatory modules across different species. Because species share common evolutionary ancestry, the core molecular machinery responsible for critical survival mechanisms is often conserved. Network alignment provides the computational methodology to identify these conserved components by mapping significant correspondences between the topologies of different species-specific networks [3–5].

However, the computational alignment of biological networks is an inherently challenging problem, mathematically related to the NP-hard Maximum Common Subgraph (MCS) problem [6, 7] This complexity is further exacerbated by the noisy, incomplete nature of inferred GRNs, which often suffer from high false-positive rates due to the underdetermined nature of transcriptomic data sets. While network alignment algorithms are well-established for Protein-Protein Interaction (PPI) networks [6, 8–10], their direct application to GRNs presents distinct algorithmic challenges. PPI alignment generally relies on global mapping strategies that seek a one-to-one node correspondence to maximize overall graph isomorphism [4, 9]. In contrast, GRNs possess a bipartite-like regulatory structure [2, 11] More importantly, evolutionary divergence, characterized by extensive species-specific gene duplication and retention events, results in many-to-many orthology mappings. Forcing a one-to-one global alignment on such data obscures the true biological architecture, necessitating local alignment strategies capable of discovering multi-node functional modules.

To bridge this methodological gap, we introduce a heuristic alignment algorithm for the cross-species discovery of conserved regulatory modules, shifting from strict topological isomorphism to a relaxed topological alignment. We formulate the discovery process as a multi-objective optimization problem that jointly maximizes sequence homology, functional coherence, and a normalized topological consensus across many-to-many ortholog tuples. Due to the exponential scaling of the search space, we implement a greedy seed-and-extend search algorithm bounded by a dynamic *ϵ*-stopping condition. This data-driven threshold dictates convergence by evaluating the marginal gain of frontier expansion, successfully preventing the functional dilution typical of unconstrained local aligners.

We validate our algorithm using high-resolution, time-series transcriptomic data from three evolutionarily distinct plant species (*Arabidopsis thaliana* [12], *Zea mays* [13], and *Sorghum bicolor* [14]) subjected to drought stress. We demonstrate that while strict topological alignment is limited to extracting small, fragmented upstream regulatory cores, our relaxed topological alignment successfully navigates the rugged search space to extract a massive, heavily connected module. By relaxing the strict intersection constraint, the algorithm recovers the dense downstream phenotypic machinery that is completely omitted by traditional methods. Overall, our algorithm provides a computational method for comparative network biology, enabling the robust projection of regulatory logic from model organisms onto complex agricultural targets.

## Materials and methods

### Problem Definition

Let *G* = *{G*_1_, …, *G*_*n*_*}* be a set of *n* GRNs from *n* distinct species. Each network *G*_*i*_ = (*V*_*i*_, *E*_*i*_) is an undirected graph representing genes (*V*_*i*_) and their regulatory interactions (*E*_*i*_).

#### Biological Prior Knowledge

To guide the alignment, we incorporate sequence homology and functional annotations.

We define an orthology set 𝒪, where each orthogroup *o* ∈ 𝒪 represents a set of genes across species that descended from a single common ancestral gene. This serves as a primary constraint for node mapping. A tuple of genes is considered a candidate for alignment only if they share membership in the same orthogroup.

Within an orthogroup, we quantify fine-grained similarity using a normalized BLAST BitScore. Let HomologyScore(*v*_*i*_, *v*_*j*_) be a function that quantifies the sequence homology between a node *u ∈ V*_*i*_ and a node *v ∈ V*_*j*_ (where *i≠ j*). To ensure the metric is symmetric and normalized, we define it as the ratio of the sum of pairwise bit scores to the sum of self-bit scores:

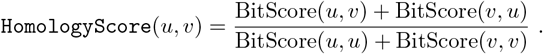

For each gene *v*, Func(*v*) denotes the set of associated Gene Ontology (GO) and KEGG pathway terms.

#### Alignment Definition

We define a *conserved subgraph alignment*, denoted as *M*, as a set of *k* disjoint node tuples:

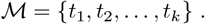

Each tuple *t*_*r*_ = (*v*_*r*,1_, *v*_*r*,2_, …, *v*_*r,n*_) represents a set of aligned homologous nodes, where *v*_*r,i*_ *∈ V*_*i*_ corresponds to a gene in species *i*. The size of the alignment is *k* = |ℳ|. Note that each tuple must contain same number of genes. All genes within a tuple *t*_*r*_ must belong to the same orthogroup *o* ∈ 𝒪. It is possible to relax this requirement. An edge is considered conserved between two tuples *t*_*p*_ and *t*_*q*_ if regulatory interactions exist between the corresponding constituent nodes in the respective networks. Specifically, it requires that (*v*_*p,i*_, *v*_*q,i*_) *∈ E*_*i*_ for all or a majority of species *i*.

For example, consider an alignment across *n* = 3 species (e.g., Species 1, 2, and 3) with size *k* = 2. The alignment ℳ = *{t*_1_, *t*_2_*}* consists of two tuples. Tuple *t*_1_ = (*v*_1,1_, *v*_1,2_, *v*_1,3_) might represent a set of homologous genes, while *t*_2_ = (*v*_2,1_, *v*_2,2_, *v*_2,3_) represents another set of homologous genes. If regulatory edges exist between these genes in all three species (e.g., (*v*_1,1_, *v*_2,1_), (*v*_1,2_, *v*_2,2_), and (*v*_1,3_, *v*_2,3_)), this structure represents a conserved regulatory subgraph.

#### Objective Function

The quality of an alignment ℳ is evaluated by an objective function *S*_total_(ℳ), calculated as the weighted sum of three biological and structural metrics. For any given pair of species *i* and *j*, these components are defined as follows.

The first component is the *homology score (S*_*homology*_), which measures the average sequence similarity across all mapped nodes:

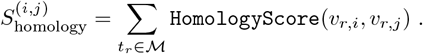

The second component is the *topology score (S*_*topology*_), which quantifies the structural coherence of the subgraphs. Rather than using a relative distance metric, this score calculates the absolute count of strictly conserved regulatory interactions mapped across the species. An edge is considered conserved if it exists between two tuples *t*_*p*_ and *t*_*q*_ in both corresponding physical networks:

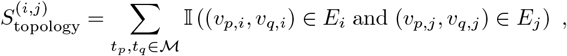

where 𝕀 (*·*) is the indicator function defined as:

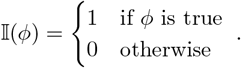

The third component is the *functional score (S*_*function*_), which measures the functional coherence among he subgraph’s target genes, calculated as the average pairwise Jaccard similarity of the annotation sets from species *i* and *j*:

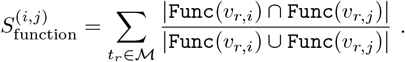

The total score *S*_total_(ℳ) of the conserved subgraph alignment ℳ is defined as the weighted sum of these pairwise components over all unique pairs of species 1 *≤ i < j ≤ n* :

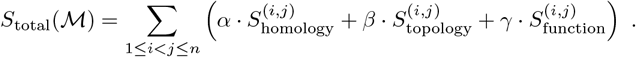

#### Complexity and Optimization

Given the set of networks 𝒢, the objective is to find a local alignment ℳ that approximates the global maximum of *S*_total_(ℳ), subject to a size constraint *k ≤ K*_max_, where *K*_max_ represents the maximum allowable size of the conserved subgraph. This constraint serves to focus the search on biologically meaningful regulatory modules rather than larger heterogeneous network components, while also ensuring computational tractability.

Since the Maximum Common Subgraph (MCS) problem is NP-hard even for *n* = 2, and the complexity increases exponentially with *n*, finding the exact global optimum for large biological networks is computationally intractable. Therefore, we approach this optimization problem using a heuristic seed-and-extend strategy. This method prioritizes local optimality at each step to construct high-scoring alignments within the bounded search space defined by *K*_max_.

### Conserved Subgraph Discovery

To identify conserved regulatory subgraphs among species, we developed a heuristic algorithm based on a greedy seed-and-extend strategy. We compare a high-stringency baseline using Strict Topological Alignment (STA) against our proposed Relaxed Topological Alignment (RTA), which is designed to discover high-scoring conserved subgraphs by iteratively growing initial seed pairs into larger subgraphs.

#### Objective Function Refinement

To evaluate a candidate alignment ℳ, we employ a multi-objective scoring function designed to reward both biological homology and structural consensus while penalizing small, fragmented subgraphs as follows:

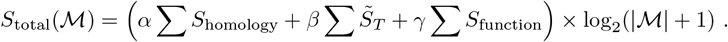

Here, 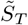 is the min-max normalized topological score, representing the normalized topological consensus. To achieve this, we first normalize the pairwise topological consensus. By dividing the raw consensus edge count 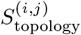 by the maximum theoretical edges between the tuples, we establish a strict [0, 1] normalization:

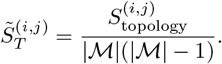

The global topological consensus is then aggregated. Let 𝒮 denote the set of species sharing the conserved edges. To prioritize evolutionary conservation, the consensus is weighted by the squared count of species sharing the edge, leading to the final macro-level transformation:

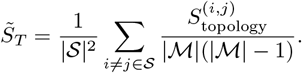

An edge is only scored if it is conserved in *m* ≥ 2 species. The raw edge count is weighted by the squared count of species sharing the edge to prioritize evolutionary conservation. The log_2_ term serves as a reward, providing the initial numerical boost required for small subgraphs to overcome the epsilon boundary.

##### Algorithm 1

Relaxed Topological Alignment with Dynamic Stopping.

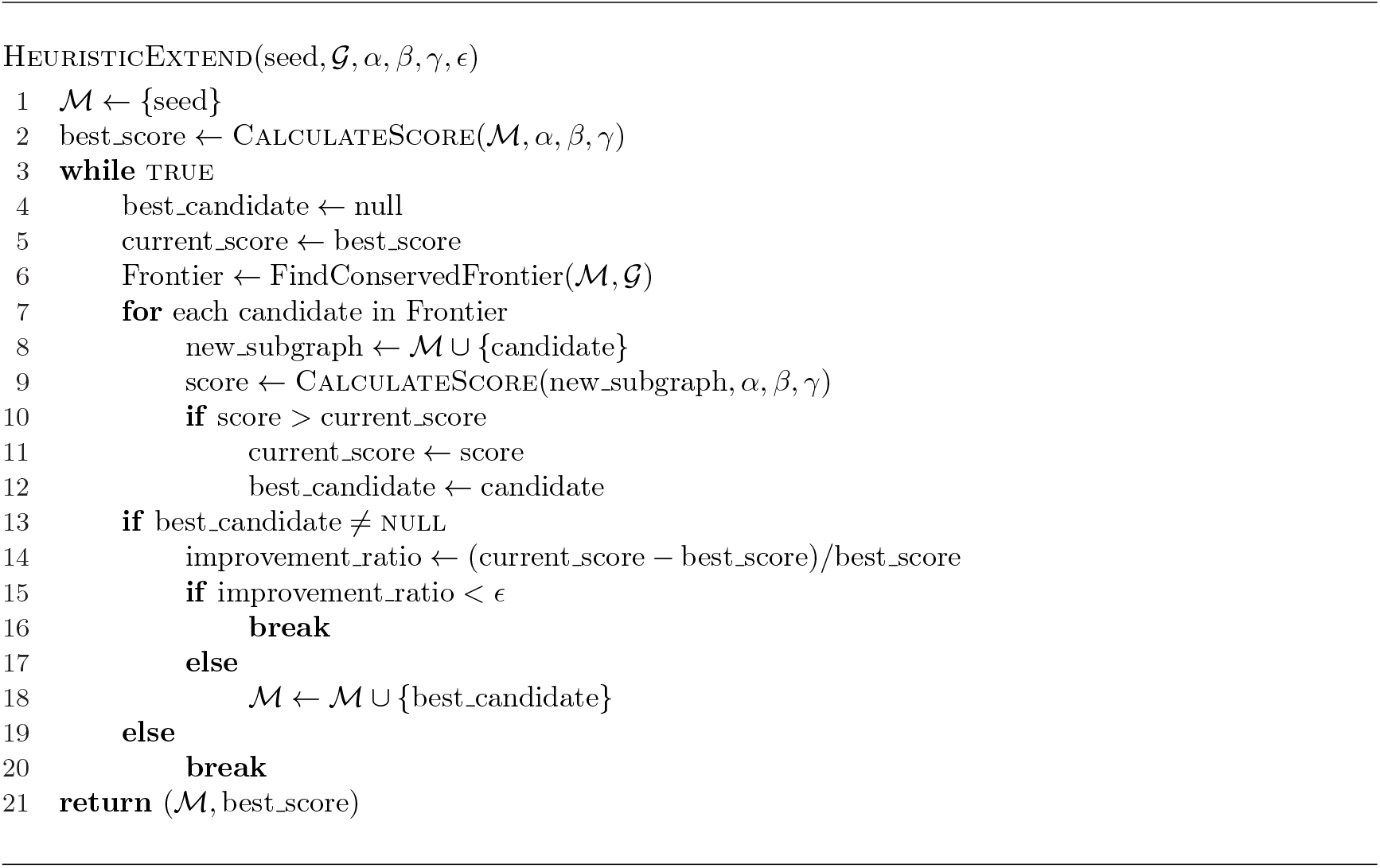

### Strict Topological Alignment

As a baseline for comparison, we define a Strict Topological Alignment (STA). The strict approach serves as a lower bound for recall. It requires perfect topological conservation. An edge is only included in the alignment if it exists across all input GRNs. While this eliminates topological false positives, it is highly susceptible to false negatives caused by genome duplications or sparse data.

### Relaxed Topological Alignment with Dynamic Stopping

Our primary contribution is a greedy seed-and-extend algorithm that discovers network boundaries with an epsilon threshold (*ϵ*). The discovery procedure is detailed in Algorithm 1 and consists of three logical phases: seed generation, greedy extension, and convergence.

#### Seed Generation

The search process is initiated from a set of high-confidence seed pairs. A seed is a homologous tuple (*v*_1_, *v*_2_, …, *v*_*n*_), where every gene belongs to the same orthogroup *o*∈𝒪. To ensure relevance, we first identify all homologous pairs that exist as nodes in both input networks. We then sort these relevant pairs in descending order based on their homology score. The algorithm iterates through the top *N* seeds, using each as a starting point for the heuristic expansion, so that the search begins in regions of high biological relevance.

#### Greedy Extension

For each seed, the algorithm performs an iterative extension to grow a larger subgraph. At each step, the algorithm identifies the Frontier (𝒞). The frontier is defined as the set of all candidate tuples *t*_*cand*_ ∉ *M* that share at least one topological edge with the current module ℳacross at least two species. To maintain computational efficiency, we only evaluate tuples that belong to orthogroups not already represented in ℳ.

We employ a greedy local search where we select the candidate *t*_*best*_ that yields the maximum value of *S*_*total*_(ℳ ∪ {*t*}).

#### Dynamic Convergence

Instead of an arbitrary size limit (*K*_max_), the expansion stops when the marginal gain in the objective function falls below a percentage of the current score. We introduce the dynamic stopping condition (Δ_*ratio*_ *< ϵ*). The algorithm evaluates the marginal gain provided by the addition of *t*_*best*_ relative to the current module score. If this ratio falls below the user-defined threshold *ϵ*, the algorithm concludes that further expansion would result in functional dilution rather than refinement, and the search terminates. This mechanism allows for the discovery of biological pathways of varying scales while ensuring the statistical significance of the conserved core.

### Data Sets

To evaluate the effectiveness of the heuristic alignment algorithm, we utilized transcriptomic and genomic data sets from three plant species: *Sorghum bicolor, Zea mays* (maize), and A*rabidopsis thaliana*.

#### Transcriptomic Data Sets

The regulatory networks were inferred from high-resolution RNA-Seq time-series data sets specifically focused on drought and developmental stress responses. For *Sorghum bicolor*, we utilized 23 samples covering four distinct time points (Day 21 to Day 42) from the BioProject PRJNA527782 [14]. The *Zea mays* data set provided a dense 32-sample time-series (Day 3 to Day 10) under the BioProject PRJNA483231 [13]. Finally, for *Arabidopsis thaliana*, we incorporated 27 samples from the BioProject PRJNA1155997 [12], which contrasts control conditions with a rigorous drought progression across six time points. The specific metadata and sample distributions for these transcriptomic resources are summarized in Table 1.

**Table 1.** Summary of transcriptomic datasets used for Gene Regulatory Network inference.

| Species | BioProject | Samples | Conditions/Time Points | Reference |
| --- | --- | --- | --- | --- |
| *S. bicolor* | PRJNA527782 | 23 | Day 21, 28, 35, 42 | [14] |
| *Z. mays* | PRJNA483231 | 32 | Day 3, 4, 5, 6, 7, 8, 9, 10 | [13] |
| *A. thaliana* | PRJNA1155997 | 27 | Control/Drought (6 points) | [12] |

#### Genomic Data Sets

To establish the homology mapping (𝒪) required for alignment, we integrated high-quality genome assemblies and annotations for each species. Genomic sequences for *S. bicolor* (JGI v3.1) and *Z. mays* (NAM v5.0) were sourced to represent the genetic complexity of cereal crops, with coding gene counts of 34,211 and 39,756, respectively [15, 16]. *A. thaliana* data was sourced from Araport11 (TAIR10) [17], representing a more compact dicot genome with 27,655 coding genes. Detailed assembly versions and chromosome counts are provided in Table 2.

**Table 2.** Genomic assembly and annotation metadata.

| Species | Source | Assembly/Annotation | Coding Genes | Chromosomes |
| --- | --- | --- | --- | --- |
| *S. bicolor* | JGI v3.1 | v3.0 / v3.1.1 | 34,211 | 10 pairs |
| *Z. mays* | NAM v5.0 | Zm-B73-NAM-5.0 | 39,756 | 10 pairs |
| *A. thaliana* | Araport11 | TAIR10 / Araport11 | 27,655 | 5 pairs |

### Homology and Orthogroup Inference

Homology scores were derived from protein sequences, as amino acid sequences are typically more conserved than nucleotide sequences [18], allowing for the detection of more distant evolutionary relationships.

All-versus-all homology searches were performed using the BLAST+ suite (v2.9.0) [19], specifically the `blastp` algorithm.

OrthoFinder [20] was subsequently applied to these proteomes to identify the orthogroups used as the fundamental units of our alignment tuples.

### Gene Regulatory Network Inference

The construction of the species-specific GRNs followed a multi-stage computational pipeline, integrating rigorous transcriptomic preprocessing with a supervised ensemble learning framework.

#### Transcriptomic Preprocessing and Quantification

Raw RNA-seq reads for all species were subjected to quality control using Trimmomatic (v0.39) [21], employing a sliding window filter (4-base window, quality cutoff of 15) and a minimum length threshold of 36 bp. Following filtering, transcript abundances were quantified using the quasi-mapping tool Kallisto (v0.46.1) [22]. To ensure cross-species comparability, transcript-level counts were collapsed to gene-level abundances (TPM) using the tximport package [23]. Potential regulators were identified by cross-referencing all coding genes against the Plant Transcription Factor Database (PlantTFDB) [24].

#### Differential Expression and Node Set Expansion

To define the initial node sets for GRN inference, we performed a differential expression (DE) analysis for each species under drought stress. We identified differentially accumulated transcripts (DATs) using a consensus approach that integrated three leading methods: DESeq2 [25], edgeR [26], and limma-voom [27].

For downstream analysis, we only considered transcripts identified as significant by all three methods. This initial consensus yielded 1,112 DATs for *Z. mays*, 1,790 for *A. thaliana*, and 329 for *S. bicolor*.

The significantly lower DATs count observed in *S. bicolor* likely stems from the lower temporal resolution of its transcriptomic sampling (4 time points) compared to *Z. mays* (8) and *A. thaliana* (6), which limits the statistical power to detect transient regulatory shifts. To mitigate this sampling bias and reduce the false negative rate, we expand the DATs set for each species based on the homology.

We leveraged the 10,429 orthogroups (𝒪) where all three species were represented. A candidate recovery rule was defined such that for any orthogroup *o* ∈𝒪, if at least two species exhibited a consensus DE response for their respective genes, all genes within that orthogroup were included in the final GRN input set.

This process identified 2,285 high-confidence orthogroups for expansion, effectively balancing the node counts across the three networks as shown in Table 3,. By using interspecies consistency as a biological filter, we ensured that the resulting GRNs were enriched for genes with a high probability of conserved regulatory function, providing a more equitable foundation for the subsequent network alignment.

**Table 3.** Node set expansion via orthogroup-informed recovery.

| Species | Initial DATs | Expanded Node Set | % Increase |
| --- | --- | --- | --- |
| *Z. mays* | 1,112 | 1,359 | 22.2% |
| *S. bicolor* | 329 | 680 | 106.7% |
| *A. thaliana* | 1,790 | 2,031 | 13.5% |

#### Two-Level Stacking Ensemble Architecture

We employed a two-level stacking ensemble to produce high-confidence regulatory maps. The framework integrates five heterogeneous base learners, each optimized for specific aspects of time-series data: SWING [28], GENIE3 [29], dynGENIE3 [30], PEAK [11], and OutPredict [31].

The confidence scores 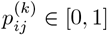from these five base methods were aggregated by a logistic regression metalearner. This metalearner was pre-trained on augmented DREAM4 *in silico* benchmarks [32–35] to learn the optimal weighting for each base learner’s predictions. To ensure the reliability of the final edge weights, the model incorporated sigmoid-based probability calibration [36] and the final interaction thresholds was optimized to maximize the F1-score.

## Results

Nulla mi mi, venenatis sed ipsum varius, Table **??** volutpat euismod diam. Proin rutrum vel massa non gravida. Quisque tempor sem et dignissim rutrum. Lorem ipsum dolor sit amet, consectetur adipiscing elit. Morbi at justo vitae nulla elementum commodo eu id massa. In vitae diam ac augue semper tincidunt eu ut eros. Fusce fringilla erat porttitor lectus cursus, vel sagittis arcu lobortis. Aliquam in enim semper, aliquam massa id, cursus neque. Praesent faucibus semper libero.

### Genomic Characterization

We first characterized the conservation landscape of the input data sets to establish the boundary of the alignment search space. OrthoFinder [20] successfully assigned 88,502 genes (87.2% of the total gene pool) to 21,692 orthogroups. The distribution of these groups indicates a high degree of core genetic conservation. We identified 10,429 orthogroups containing at least one representative from all three species, 3,973 of which consist entirely of single-copy genes.

To quantify the density of this sequence clustering, we calculated the G50 and O50 statistics. The G50 metric represents the orthogroup size at which 50% of all assigned genes reside in orthogroups of that size or larger, while O50 indicates the minimum number of orthogroups needed to encompass half of all assigned genes. Our analysis yielded a G50 of 4 and an O50 of 7,556, suggesting a balanced distribution of gene family sizes across the selected species.

### Inferred GRNs and Orthology Mapping

To establish the basis for cross-species alignment, we first characterized the topological dimensions of the individual GRNs and their respective orthology overlaps. To eliminate computational noise and focus strictly on high-confidence regulatory interactions, the raw ensemble GRN output for each species was pruned to retain only the top 10,000 edges ranked by their consensus weights. This cutoff resulted in highly sparse adjacency matrices (densities ranging from 0.002 to 0.023), ensuring that any topological consensus identified during cross-species alignment represents strong, evolutionarily conserved regulatory signals.

As summarized in Table 4, the inferred networks exhibit varying degrees of structural complexity due to their different vertex set sizes. *A. thaliana* exhibited the highest connectivity with 2,017 nodes (including 155 TFs) with an average degree of 4.96, yielding the lowest graph density (0.0025). *Z. mays* and *S. bicolor* contained 1,352 nodes (average degree 7.40) and 657 nodes (average degree 15.22), respectively. A total of 141 orthogroups (OGs) were found to be common across all three species’ inferred networks, representing the potential core regulome available for strict three-way alignment.

**Table 4.** Summary of individual species GRNs and orthology representation.

| Metric | *A. thaliana* | *Z. mays* | *S. bicolor* |
| --- | --- | --- | --- |
| Total Nodes (Genes) | 2,017 | 1,352 | 657 |
| Total Transcription Factors | 155 | 108 | 42 |
| Average Degree | 4.96 | 7.40 | 15.22 |
| Network Density | 0.0025 | 0.0055 | 0.0232 |
| Represented Orthogroups | 1,367 | 863 | 365 |
| Non-orthologous Genes | 230 | 149 | 27 |

### Baseline Performance of Strict Topological Alignment

To establish a comparative baseline, we first applied the STA detailed in the method section to identify the most robustly conserved regulatory interactions across the three species. By performing an exhaustive search of the 141 core orthogroups, we generated a search space of 5,639 potential multi-dimensional gene tuples. Under this strict regime, an edge is only retained if it exists in the inferred GRNs of all three species simultaneously. This stringent mathematical intersection effectively filtered out 99.9% of species-specific noise, yielding only 104 strictly conserved edges.

From this filtered interactome, we identified 10 non-trivial conserved structures, encompassing a total of 75 tuples and 104 inter-tuple edges. Here, we define a conserved module as a connected component within the alignment graph containing at least two disjoint tuples joined by at least one strictly conserved inter-tuple edge. The largest discovered module consists of 51 tuples and 87 edges. To visualize the physical regulatory wiring of this specific sub-structure, we projected it back into species-specific networks as shown in Figure 1. As illustrated, we observed a diverse regulatory composition: the *Z. mays* component includes 14 genes (7 TFs), the *S. bicolor* component includes 17 genes (7 TFs), and the *A. thaliana* component includes 11 genes (6 TFs). This suggests that while the edges are strictly conserved, the underlying gene duplication and retention patterns within each species provide a unique topological structure for the same functional unit.

To assess the functional coherence of this strictly conserved module, we conducted a Gene Ontology (GO) enrichment analysis on the *A. thaliana* gene members. The analysis returned 32 significantly enriched terms (FDR *<* 0.05), primarily centered on abiotic stress response and photosynthetic regulation. Notably, we observed a high enrichment of terms related to the cellular response to water deprivation (GO:0042631) and response to desiccation (GO:0009269), alongside specific pathways for abscisic acid (ABA) stimulus (GO:0071215) and regulation of stomatal closure (GO:0090333).

**Fig 1.**
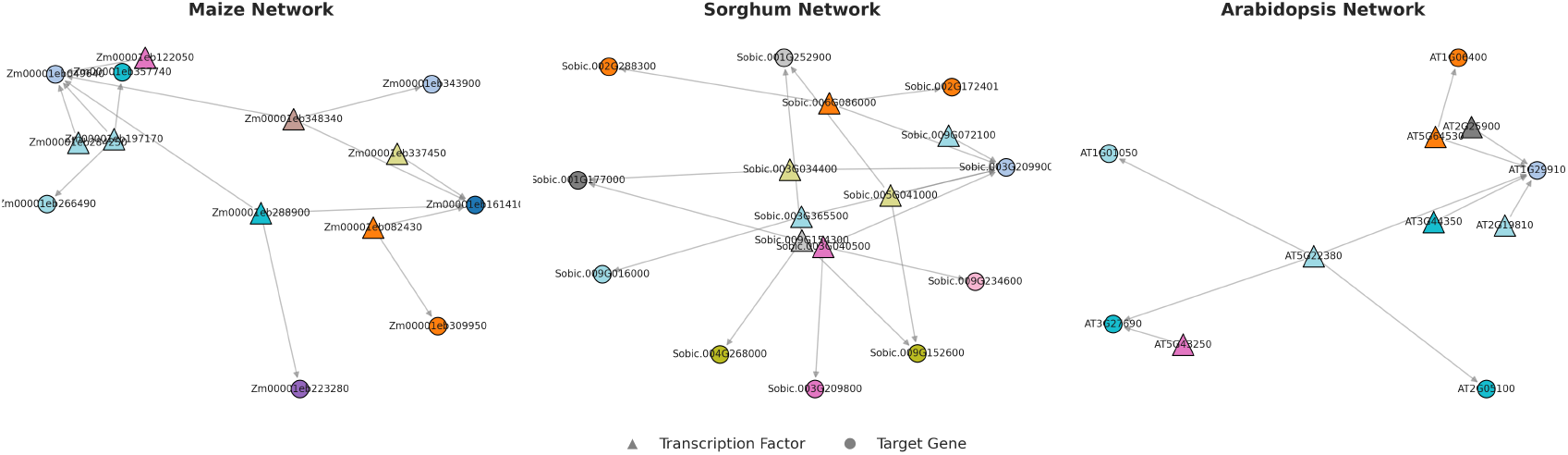
Species-specific topological projections of the largest strictly conserved regulatory module. The cross-species alignment module (comprising 51 tuples) was mapped back to the individual interactomes of *Zea mays, Sorghum bicolor*, and *Arabidopsis thaliana*. Triangles represent known transcription factors (TFs), while circles denote downstream target genes. Node colors indicate cross-species orthogroups, with identical colors across the three panels representing orthologous genes. The projected subnetworks highlight species-specific topological variations stemming from evolutionary duplication and retention, consisting of 14 nodes and 14 edges in *Z. mays*, 17 nodes and 18 edges in *S. bicolor*, and 11 nodes and 10 edges in *A. thaliana*.

Furthermore, the module was also enriched for terms related to the maintenance of the photosynthetic apparatus, including Photosystem I and II (GO:0009522, GO:0009523) and chlorophyll binding (GO:0016168). The presence of positive regulation of reactive oxygen species (GO:1903428) might suggest that this conserved core represents a coordinated regulatory module.

Subsequent isolation of *A. thaliana* genes within this module associated with DNA-binding transcription factor activity (GO:0003700) identified specific node-to-node correspondences for five key transcription factor families, including members of *TZF1/2* RNA-turnover regulators, *NAC* family transcriptional repressors, and *XND1*. In particular, *XND1* is a specialized TF in the NAC-domain that was previously implicated in the regulation of drought responses and the coupling of root xylem plasticity with *Na*^+^ unloading [37, 38].\ Detailed ortholog mappings are provided in Table 5.

**Table 5.** Detailed orthologous mapping of core regulatory TFs across the three-species alignment.

| *A. thaliana* TF | *Z. mays* Ortholog(s) | *S. bicolor* Ortholog(s) |
| --- | --- | --- |
| ***OZF1/TZF2***<br>(AT2G19810) | Zm00001eb284250,<br>Zm00001eb337450 | Sobic.009G072100,<br>Sobic.003G034400 |
| ***ATCTH/TZF1***<br>(AT2G25900) | Zm00001eb284250,<br>Zm00001eb337450 | Sobic.003G034400,<br>Sobic.009G072100 |
| ***NAC061***<br>(AT3G44350) | Zm00001eb288900,<br>Zm00001eb197170,<br>Zm00001eb348340 | Sobic.005G041000,<br>Sobic.003G365500 |
| ***NAC090***<br>(AT5G22380) | Zm00001eb288900,<br>Zm00001eb197170,<br>Zm00001eb348340 | Sobic.009G154300,<br>Sobic.003G365500,<br>Sobic.005G041000 |
| ***NF-YC13*** (AT5G43250) | Zm00001eb122050 | Sobic.003G040500 |
| ***XND1*** (AT5G64530) | Zm00001eb082430 | Sobic.006G086000 |

The concurrent structural alignment of these transcription factors confirms the isolation of a complex regulatory subnetwork. By mapping the topological dependencies between *NAC* repressors and xylem modulators like *XND1*, the algorithm successfully extracts a shared network architecture deployed during extreme drought stress across evolutionarily distinct species.

To further validate the functional relevance of the strictly aligned core, we analyzed the temporal expression trajectories of a sub-module comprising 11 conserved tuples. These tuples were selected based on their high co-annotation for drought-related metadata, including response to desiccation (GO:0009269), response to water deprivation (GO:0042631), and response to abscisic acid (GO:0071215). The sub-module contains specific orthologous nodes belonging to the Light-Harvesting Chlorophyll B-binding (*LHCB*) family, including *LHCB2*.*1* (AT2G05100) and *LHCB2*.*3/2*.*4* (AT3G27690) in *A. thaliana*, with corresponding orthologs in *Z. mays* and *S. bicolor* (Table 6).

**Table 6.** Accession numbers for orthologous nodes within the LHCB-enriched sub-module.

| Species | Gene Accession (Node ID) | Functional Symbol |
| --- | --- | --- |
| *A. thaliana* | AT2G05100 | *LHCB2.1* |
|  | AT3G27690 | LHCB2.3/DEG13 |
| *Z. mays* | Zm00001eb049640 | *lhcb9* |
|  | Zm00001eb161410 | — |
|  | Zm00001eb343900 | *lhcb3* |
|  | Zm00001eb357740 | *lhcb10* |
| *S. bicolor* | Sobic.001G177000 | — |
|  | Sobic.003G209800 | — |
|  | Sobic.003G209900 | — |
|  | Sobic.009G234600 | — |

We computed *Z*-score normalized expression profiles for these member genes across the drought time-series to evaluate temporal signal divergence. As illustrated in Figure 2, the temporal dynamics exhibit significant variance, indicating that the conserved topology operates under different systemic latency and activation thresholds depending on the host system. The *A. thaliana* genes exhibits an immediate decrease upon stress onset. However, it displays a secondary transcriptomic surge during the Day 20–27 interval before a final decline at Days 27–29, reflecting a putative strategy to prioritize survival and reproduction. This complex pattern is supported by high-resolution time-course transcriptomics demonstrating that the drought response in this annual species operates through multiple overlaying transcriptional programs, with gradual onset responses heavily overlapping with the plant aging program [12]. Specifically, *A. thaliana* undergoes a transcriptional rejuvenation during sublethal drought before returning to its baseline aging trajectory, yielding the secondary amplitude surge observed in our network projections. Conversely, the responses of the C4 species reflects their higher drought tolerance thresholds. *Z. mays* shows a transient activation signal around Days 3–4, followed by a sustained decrease phase. *S. bicolor* maintains a delayed and slight elevation in expression during its early prolonged stress phase (Days 21–28) before eventually decreasing as the drought stress persists. This algorithmic extraction of the *LHCB* temporal sub-module perfectly aligns with field-based transcriptomic and proteomic validations, which observed significant down-regulation of these exact *S. bicolor* photosystem II *LHCB* homologs under postflowering drought stress [14]. Similarly, transcriptomic profiling in *Z. mays* confirms that severe perturbations of photosynthesis-related genes primarily manifest at the late drought stage [13]. This may imply that the underlying execution of this conserved module is finely modulated by the species-specific tolerance ability. Crucially, this result highlights that our strict alignment algorithm can successfully identify the conserved structural core (e.g., ABA-target interactions) even when the actual temporal expression and activation thresholds vary between species.

**Fig 2.**
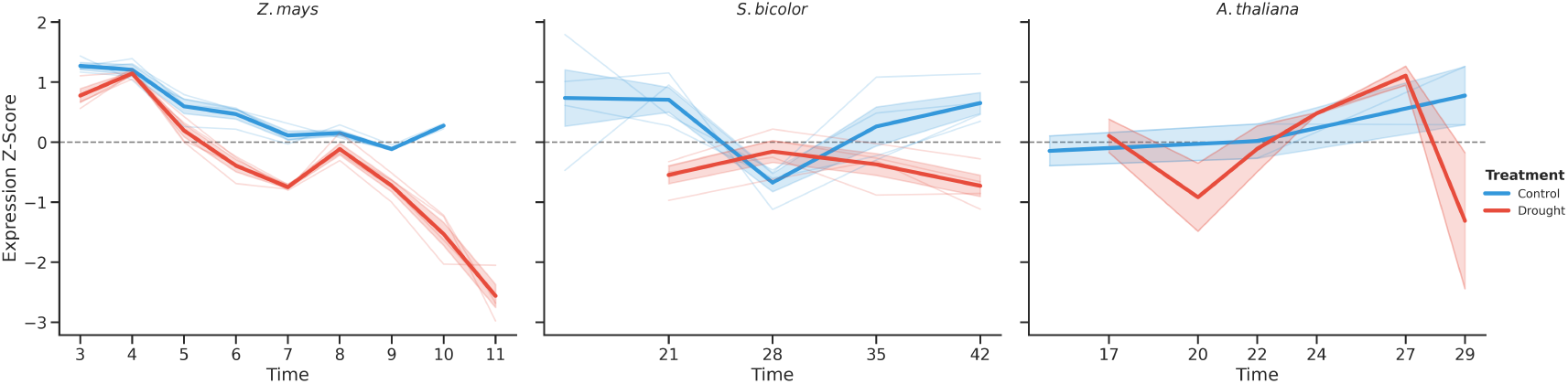
Conserved temporal expression dynamics of the core drought-responsive module. *Z*-score normalized expression trajectories are shown for the 11 conserved orthologous tuples across *Z. mays, S. bicolor*, and *A. thaliana*. Faint lines trace the expression profiles of individual genes, while bold lines represent the module-wide average expression, with shaded regions indicating the standard error. Under drought stress, the C4 crops (*Z. mays* and *S. bicolor*) exhibit a synchronized down-regulation relative to well-watered controls. In contrast, the C3 species (*A. thaliana*) displays a divergent regulatory signal, maintaining elevated expression levels during the late-stage drought interval (Days 20–27).

### Relaxed Topological Alignment Performance

We evaluated the performance of the relaxed topological alignment using a weighted configuration of *β* = 0.6 (topology) and *γ* = 0.4 (function). The search was initialized with *N* = 10 independent seeds and a marginal improvement threshold of *ϵ* = 0.01. As shown in the expansion trajectories (Figure 3), the modules exhibited a rapid growth before reaching the *ϵ*-driven convergence point. Module sizes ranged from 1 to 173 tuples, demonstrating that the dynamic stopping condition successfully bounds the graph expansion and prevents functional dilution in larger components.

**Fig 3.**
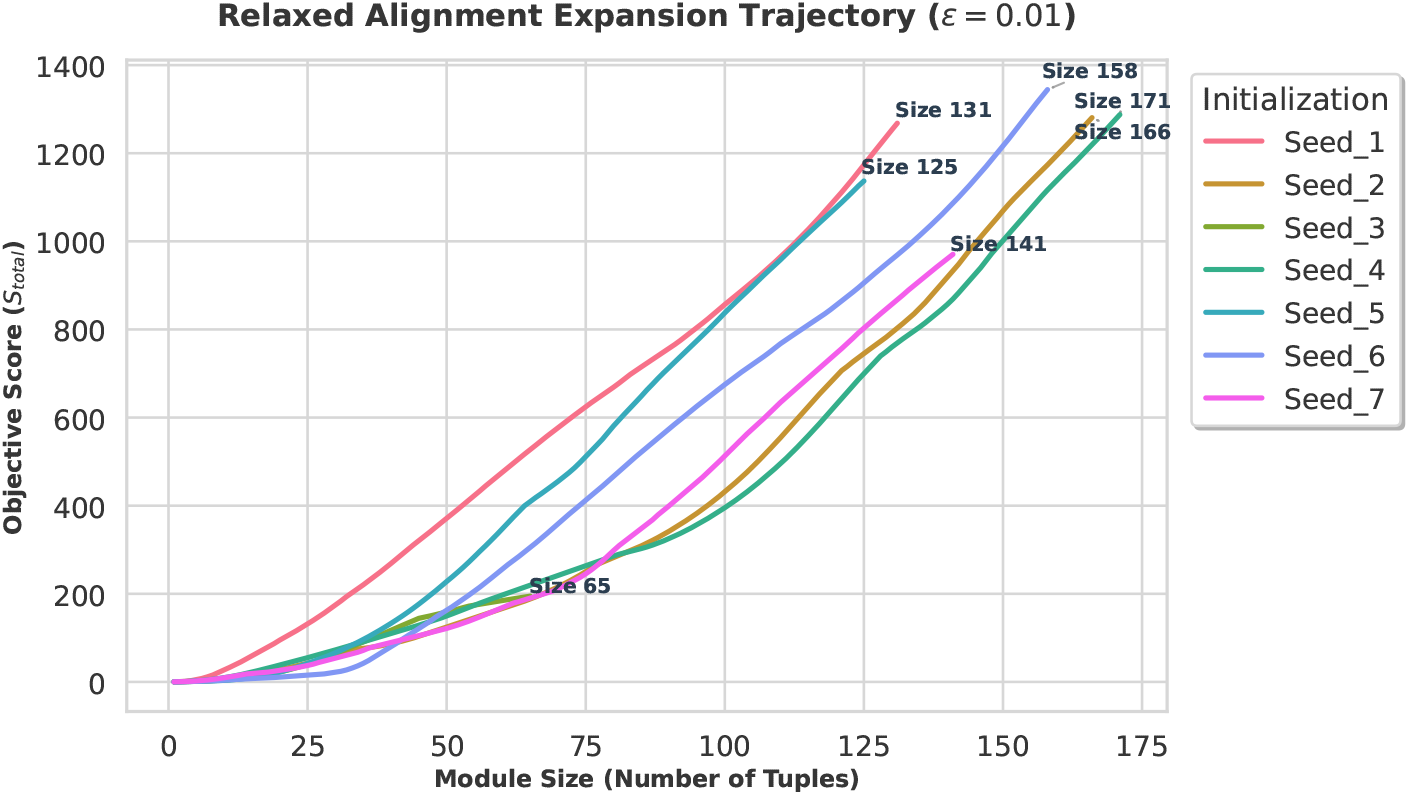
Growth trajectories of the relaxed topological alignment. The line plot traces the step-wise heuristic expansion of the conserved module originating from distinct valid initial seeds. The x-axis represents the expansion step, which corresponds to the cumulative module size (number of tuples). The y-axis tracks the corresponding global objective score (*S*_*total*_). The expansion iteratively recruits neighboring tuples that maximize the objective score. To prevent unconstrained propagation into topological noise, the algorithm employs an early-stopping criterion, terminating the growth when the marginal score improvement drops below the predefined threshold (*ϵ* = 0.01). The convergence of these distinct trajectories toward massive, stable module sizes (e.g., sizes *>* 130) visually demonstrates the algorithmic robustness and the clear boundary of the underlying biological network.

The 10 discovered modules revealed high levels of topological overlap. Despite being initialized from diverse, non-adjacent seeds, the expansion process resulted in significant overlap between independent runs. After pruning isolated seeds that failed to recruit additional nodes (size *k* = 1), we computed the intersection of the effectively expanded modules, visualized via an UpSet plot (Figure 4). For instance, the subgraph expanded from Seed 1 (*k* = 131) was a complete subset of the subgraph expanded from Seed 6 (*k* = 158). This substantial structural overlap across distinct initialization points demonstrates that the greedy heuristic consistently converges toward a stable, global module representing the core regulatory response, rather than diverging into disparate topological components.

**Fig 4.**
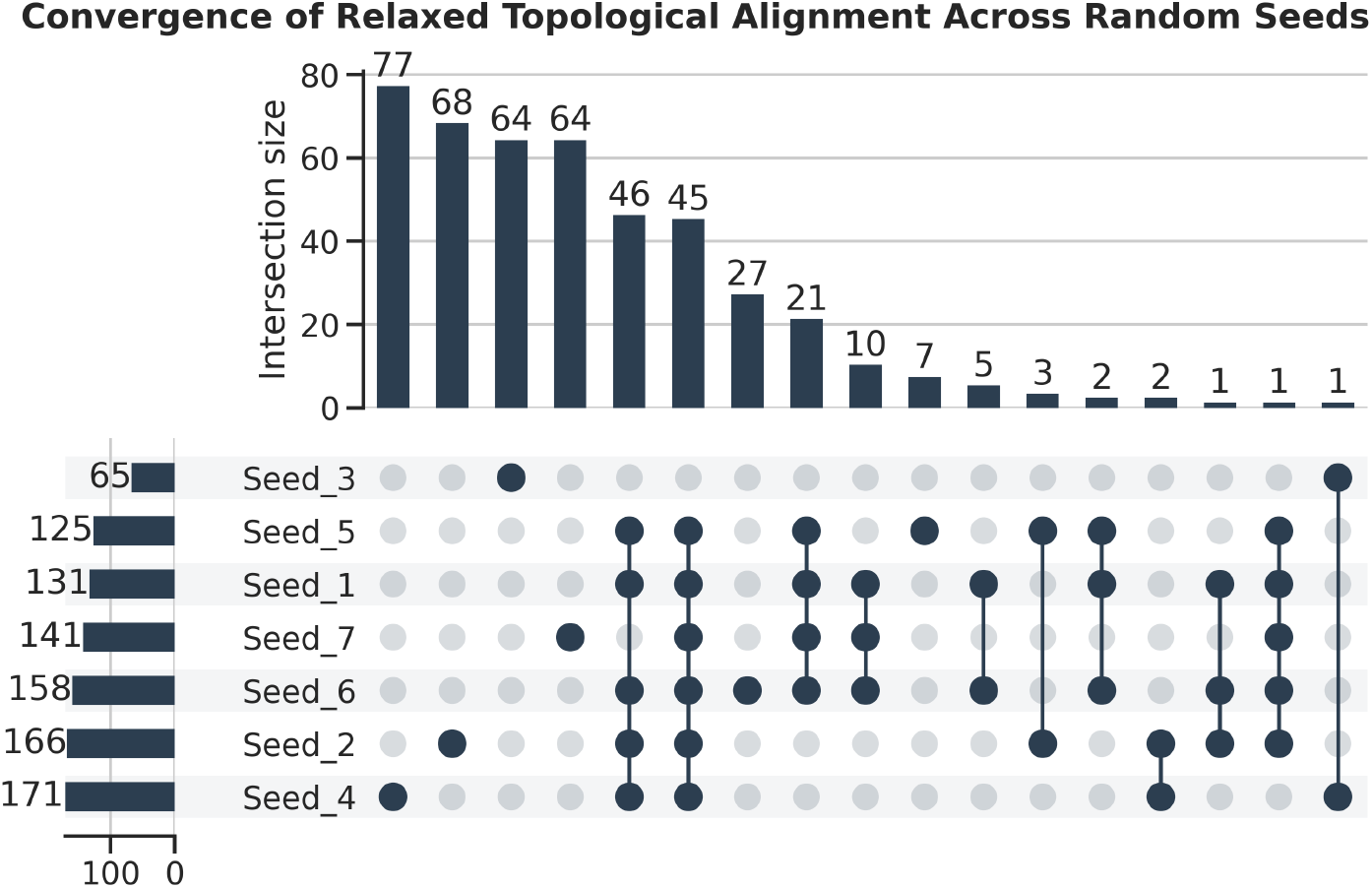
UpSet plot detailing the topological convergence of the relaxed topological Alignment. The expansion algorithm was initiated from 10 randomly selected seed tuples, excluding isolated seeds that yielded no topological expansion (module size = 1). The horizontal bar chart (bottom left) displays the total number of tuples captured by each successfully expanded seed. The vertical bar chart (top) quantifies the size of the intersections between different expanded modules, with the specific subset combinations indicated by connected solid dots in the matrix below. The substantial intersection sizes, including complete subset relationships (e.g., Seed 1 being entirely encapsulated within Seed 6), highlight algorithmic robustness and confirm that the relaxed alignment converges toward a globally stable conserved module.

To characterize the global macroscopic structure, we computed the union of all discovered modules, yielding a set of 447 unique tuples. Within this union space, we identified a single dominant connected component containing 444 tuples (99.3% of the total union). This component maps to 68 *Z. mays*, 61 *S. bicolor*, and 106 *A. thaliana* unique genes. This heavy-tailed component distribution confirms that the heuristic algorithm successfully navigates toward a primary conserved regulatory core within the input GRNs, circumventing structural fragmentation.

To examine the physical regulatory architecture of this large conserved module, we projected the 444 tuples back onto the species-specific interactomes (Figure 5). Compared to the strict baseline, the relaxed module exhibits a substantially expanded edge density while maintaining a consistent core of regulatory TFs. Specifically, the subnetwork comprises 56 nodes (15 TFs) with 96 edges in *Z. mays*, 61 nodes (15 TFs) with 242 edges in *S. bicolor*, and 57 nodes (14 TFs) with 46 edges in *A. thaliana*. This intra-species structural variance highlights how strictly conserved upstream TFs orchestrate a divergent array of species-specific downstream target topologies.

**Fig 5.**
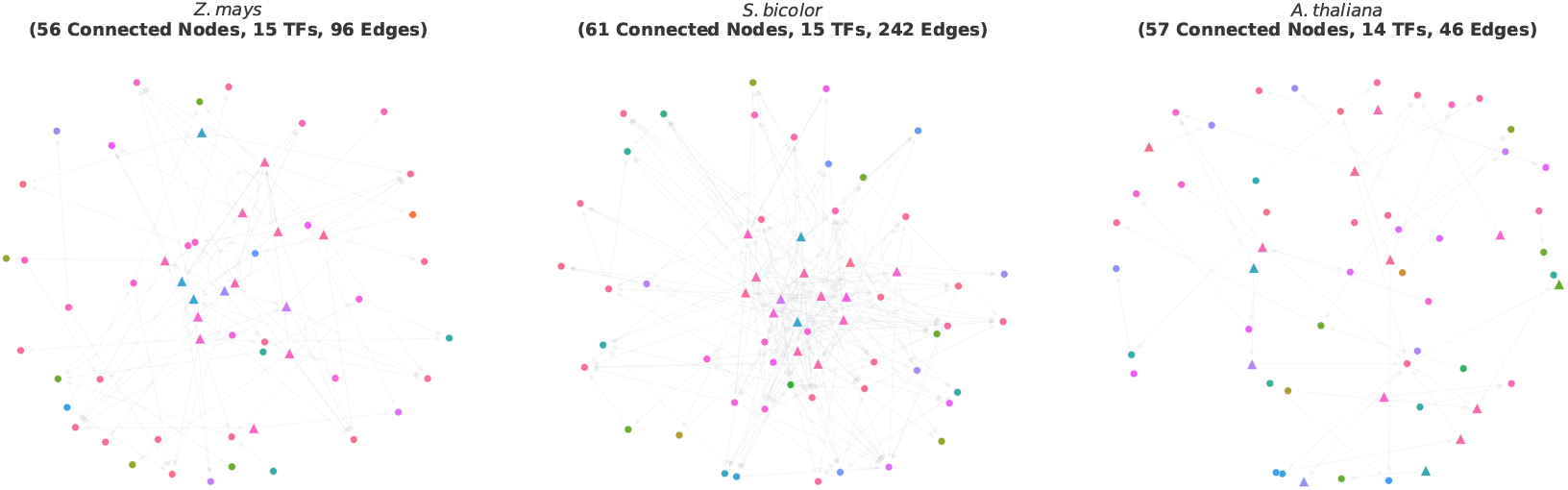
Topological architecture of the relaxed alignment result. Species-specific regulatory subnetworks of the 444-tuple Module in *Z. mays, S. bicolor*, and *A. thaliana*. Nodes are colored by their cross-species orthogroups. Triangles denote transcription factors (TFs), while circles represent downstream target genes. The relaxed alignment successfully recovers dense species-specific topological structure (e.g., 242 edges in *S. bicolor*) surrounding a conserved TF core (14-15 TFs per species).

We then performed GO enrichment analysis on the *A. thaliana* node set to validate the functional coherence of this expanded network boundary. The analysis yielded 109 statistically significant terms (FDR *<* 0.05). Specifically, the top enriched pathways alignes well with the drought response. This includes extensive cellular restructuring, including structural constituent of cytoskeleton, microtubule cytoskeleton organization, water channel activity, and water transport. The recovery of these pathways confirms that the relaxed objective function successfully links conserved upstream regulatory signals to their physiological phenotypes.

To quantify the sensitivity of the heuristic approach, we measured the recall of the 51 tuples identified by the strict baseline alignment. The heuristic result successfully recovered 46 of the 51 strict tuples, representing a recall of 90.20%. Only 5 tuples from the strict baseline were omitted from the heuristic expansion. A post-hoc analysis indicates these tuples lacked the high topological and/or functional similarity score required to surpass the *ϵ* = 0.01 threshold relative to the more dense neighborhoods discovered by the heuristic search. This indicates that our algorithm effectively balances the retention of high-precision core interactions with the expansion of high-recall functional neighborhoods.

Finally, we executed a grid search across the (*β, γ*) parameter space to evaluate the optimization trade-off between alignment size and functional purity (Figure 6). We fixed the homology coefficient *α* during this grid search to establish a stable mapping coordinate space, as sequence homology serves as a deterministic physical ground truth derived directly from observable protein sequences. Conversely, the topological edges are probabilistically inferred and subject to thresholding noise, while the functional annotations (*S*_function_) are notoriously sparse. By anchoring the search with a fixed *α*, the grid search strictly isolates the algorithmic trade-off between enforcing structural isomorphism (*β*) and leveraging incomplete semantic metadata as a heuristic guide (*γ*). As expected, the mean functional score (*S*_function_) scaled monotonically with the *γ* coefficient, demonstrating that a high *γ* allows the heuristic to bridge topological gaps using functional metadata to expand the module boundaries. Conversely, module size exhibited a non-linear response to the topological weight (*β*), peaking at *β* = 0.2 and oscillating around a mean cardinality of 130 tuples, confirming that a excessively high *β* restricts the search to exact structural matches and severely limits recall. The parameter behavior demonstrates that neither metric is strictly superior. Topological constraints define the structural boundary, while the functional objective component acts as a critical regularizer, preventing the greedy heuristic from unbounded expansion into low-confidence graph regions where inference noise dominates.

**Fig 6.**
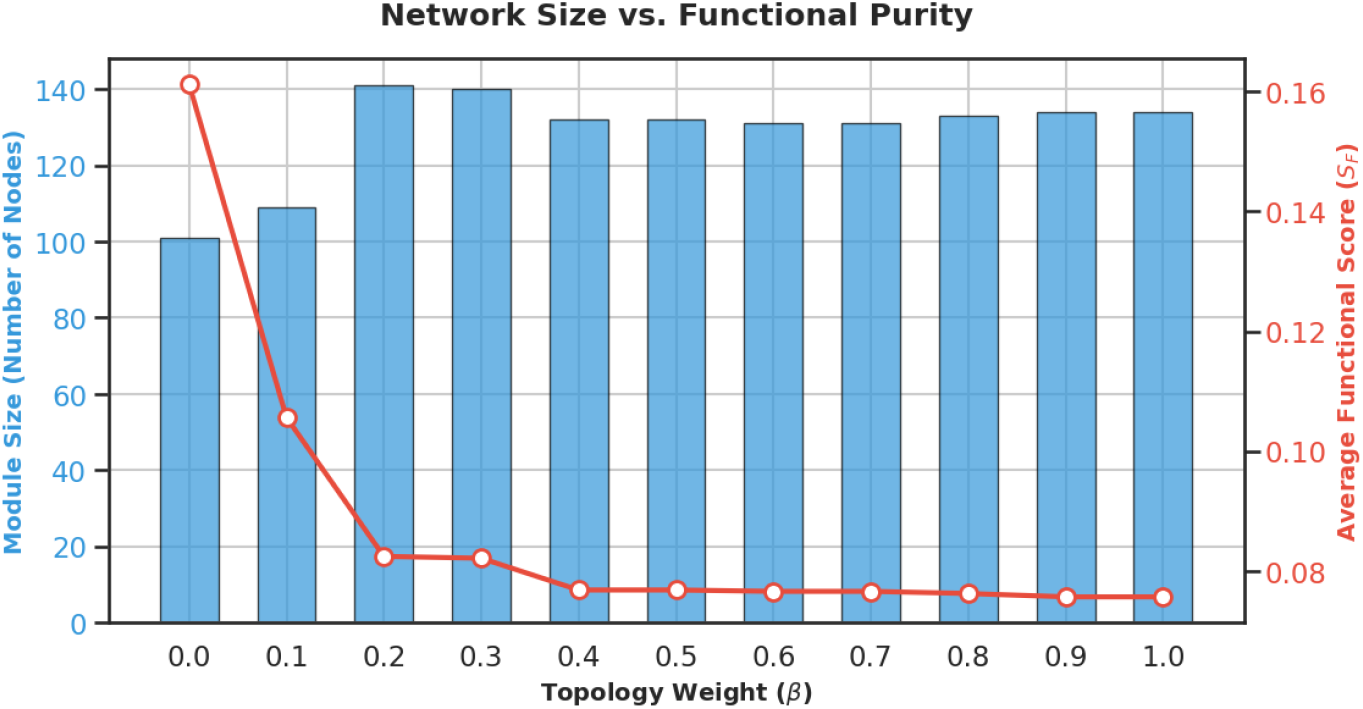
Parameter optimization for the relaxed topological alignment via grid search. An extensive grid search was performed to determine the optimal balance between the topology weight (*β*) and the functional weight (*γ*). The blue bars represent the resulting module size (number of aligned tuples) under various parameter combinations, exhibiting a distinct peak at *β* = 0.2 before reaching a plateau. The red line denotes the average functional score, which decreases monotonically as the algorithm shifts weight toward biological constraints. This non-linear relationship demonstrates that functional information acts as a critical regularizer. Integrating homology, topology, and functional metrics ensures the extraction of a network that is both topologically robust and biologically meaningful, avoiding the extremes of being overly restrictive or dominated by topological noise.

## Discussion

The recovery of the *NAC90* -*NAC61, TZF1/2*, and *XND1* nodes at the center of the alignment serves as a critical benchmark for the performance of our algorithm. By identifying these specific subgraphs through a high-stringency topological search, we demonstrate that the proposed alignment algorithm is capable of extracting coherent functional units from high-dimensional, noisy transcriptomic expression data.

This result validates the signal within the inferred GRNs. While individual GRN inference methods often suffer from high false-positive rates, the intersection of three independently inferred networks across species acts as a denoising filter. The necessity of this cross-species intersection is underscored by the massive scale of single-species stress responses. For example, field-grown *S. bicolor* exhibits significant differential expression in approximately 44% of its transcribed genome during drought [14]. When such a large proportion of the genome shifts in response to a systemic perturbation, standard single-network motif discovery is easily overwhelmed by false positives. The survival of the *NAC, XND1*, and *TZF* nodes through this filter suggests that these regulatory interactions are stable, conserved features of the stress-response architecture. This is directly corroborated by transcriptomic studies in *Z. mays*, which identify the *ZmNAC* transcription factor family as significantly enriched during the early and middle stages of progressive drought stress [13]. By extracting this specific module, our strict alignment successfully isolates the true, early-acting signaling patterns that initiate the adaptive response.

The relaxed topological alignment exclusively unveils the downstream phenotypic machinery. While the strict alignment reliably captured the conserved upstream regulatory core, it lacked the topological flexibility to evaluate the highly duplicated downstream effectors. The strict one-to-one mapping struggles to make accurate matches due to the crop’s genomic adaptation. Early and middle drought-responsive genes in diverse *Z. mays* lines display high copy number variation, including presence and absence variation, as an evolutionary mechanism to minimize fitness costs during environmental stress [13]. Functional enrichment of the relaxed alignment result revealed 83 additional GO terms entirely omitted by the strict approach. These newly captured terms encompass the definitive physical mechanisms of system adaptation, including extensive cellular structural remodeling, hydraulic regulation, and protein protection mechanisms. Our relaxed alignment successfully captures the extensive abscisic acid (*ABA*) stimulus pathways, which reflects the complex phytohormone crosstalk driving this physical execution. Progressive drought in *Z. mays* triggers a massive accumulation of *ABA*, which simultaneously orchestrates stomatal closure and negatively regulates the expression of jasmonic acid and salicylic acid pathway genes [13]. Similarly, *S. bicolor* exhibits strong up-regulation of genes involved in *ABA* catabolism and signaling inhibition to buffer the hormonal response [14]. This is further confirmed by *A. thaliana* studies, which exhibit rapid initial upregulation of *ABA* biosynthetic enzymes followed by the late-stage activation of fine-tuning enzymes [12]. The exclusive recovery of direct phenotypic markers decisively validates the algorithmic necessity of the relaxed alignment. It demonstrates that while the regulatory command center remains strictly conserved across species, the downstream functional execution is wired through a dense, flexible topology that can only be mapped by accommodating evolutionary divergence.

Furthermore, the successful alignment across *A. thaliana, Z. mays*, and *S. bicolor* highlights the algorithmic capacity of the algorithm to bridge significant evolutionary distances and distinct systemic outputs. While the continuous temporal expression analysis revealed divergent response trajectories, specifically the rapid response in the C3 model organism compared to the prolonged activation states in the C4 species, the alignment result suggest that these distinct phenotypic patterns are orchestrated by a deeply conserved topological core. This demonstrates that our algorithm can effectively project well-characterized regulatory interactions from the model organisms onto the high-dimensional target spaces of polyploid crop genomes. This capability provides a putative road map for identifying targeted genetic modifications to optimize agricultural resilience.

While the primary objective of this algorithm is the extraction of conserved regulatory cores, the aligned architecture inherently facilitates the quantification of network rewiring. By defining the conserved module as the graph intersection of regulatory edges across homologous tuples, the structural complement of this consensus represents evolutionary divergence. Within the boundaries of the 444-tuple module, the unaligned edges explicitly map the species-specific rewiring of downstream target genes. This allows for the direct extraction of divergent topologies anchored to a conserved regulatory command center. Additionally, the successful retention of the tripartite alignment demonstrates the algorithm’s robustness to phylogenetic bias. Because *Z. mays* and *S. bicolor* share a much closer evolutionary distance compared to the eudicot *A. thaliana*, their pairwise structural and homologous similarities are naturally heavily weighted. The algorithm’s ability to maintain the *A. thaliana* mapping without collapsing into a bipartite grass-specific subnetwork validates that the multi-objective normalization and dynamic convergence criteria effectively bridge massive evolutionary distances.

The strict topological alignment relies on a straightforward Boolean intersection of edge sets, operating with a low computational footprint bounded primarily by the size of the initial ortholog mapping. While this yields a highly optimized execution time, it suffers from severe topological fragmentation, returning small, disjointed subnetworks (maximum size of 51 tuples) that fail to represent the complete functional module. Conversely, the relaxed topological alignment, driven by a greedy seed-and-extend heuristic, requires iterative recalculation of the objective function across the expanding graph frontier. The worst-case asymptotic time complexity for this heuristic is bounded approximately by *O*(*N·*| *V*| ^3^), where *N* is the number of initial seeds and |*V*| is the maximum vertex set size. Although this imposes a substantially higher computational cost and longer execution times, the dynamic *ϵ*-stopping condition effectively truncates the empirical search tree, preventing an exponential explosion. The structural payoff is the successful extraction of a cohesive, 444-tuple large module that the strict approach could not assemble. Thus, the increased computational overhead of the relaxed heuristic is justified by its high recall rate and its ability to construct a connected topological subgraph from conserved regulatory cores to species-specific phenotypic outputs.

The evaluation of the relaxed topological alignment algorithm validates its capacity to extract conserved regulatory modules across highly divergent genomes. Within the strict parameters of this objective, the homology mapping and algorithmic alignment were constrained exclusively to *Arabidopsis thaliana, Zea mays*, and *Sorghum bicolor*. This constraint was implemented to test the computational framework rather than to execute an exhaustive query of the entire plant kingdom. However, because *Arabidopsis* is a dicotyledon and the latter two species are monocotyledons, their evolutionary divergence spans over 150 million years. A regulatory topology that remains strictly conserved across this massive evolutionary distance represents an ancient and fundamental survival architecture for angiosperms. This evolutionary persistence is corroborated by the network findings established in the second objective. Many of the transcription regulators driving this core module, including the *NAC* and *Trihelix* families, are the exact homologous families identified as master regulators in the resurrection plant *Selaginella sellowii*.

Consequently, the extracted 444-tuple module serves as a highly validated regulatory blueprint. Future research does not require the computationally expensive reconstruction of global networks for every newly sequenced genome. Instead, algorithms can utilize this conserved module as a direct query subgraph against genomic databases, such as Phytozome, to determine its presence in other agricultural crops or nonvascular plants. This targeted projection will allow researchers to mathematically trace the exact evolutionary assembly point of this stress response engine.

The success of this module extraction relies heavily on the specific weighting of the optimization function with multiple objectives. Within this mathematical formulation, the sequence homology score acts as the most critical parameter. While topological consensus and functional coherence serve as necessary regularizers, sequence homology anchors the exponentially scaling search space to biological reality. Without a stringent sequence homology constraint, a relaxed topology metric would permit the algorithm to align functionally unrelated subgraphs that merely possess similar degree distributions. By prioritizing sequence homology, the algorithm restricts the heuristic search and ensures that the mapped graphs represent true orthologous genetic lineages rather than stochastic structural noise or isomorphic graph artifacts.

Furthermore, the mathematical architecture of this alignment algorithm is completely agnostic to species boundaries. The framework can be directly deployed to align regulatory networks extracted from different physical tissues within a single organism. In such an application, the sequence homology term could be updated with a paralogy mapping metric or an expression correlation score, while the functional and topological optimization functions remain mathematically identical. This adaptation would enable the computational quantification of how a singular genetic circuit undergoes structural rewiring to execute specialized physiological functions across distinct cellular environments.

## Conclusion

In this study, we formalized and addressed the challenge of discovering conserved regulatory modules across species using inherently noisy, inferred GRNs. First, we define a new network alignment problem specifically tailored for GRNs, addressing a structural challenge that has not been recognized before. Second, we provide two greedy algorithms to solve the problem for two or more GRNs from diverse species at once. Third, we test our two algorithms on drought treated RNA-seq data from three well-studied plant species.

By shifting from a paradigm of strict topological alignment to a relaxed topological alignment, our greedy algorithm successfully balances topological precision with biological recall. The implementation of an *ϵ*-bounded dynamic stopping condition effectively controls the asymptotic search space expansion while avoiding the functional dilution characteristic of large, heterogeneous graph components. Empirical validation across *A. thaliana, Z. mays*, and *S. bicolor* demonstrated that while strict alignment isolates high-confidence upstream regulatory centers, it fails to traverse the highly duplicated downstream effector networks. Our relaxed objective function successfully mapped the dense topological wiring that connects conserved transcriptional regulators to their species-specific phenotypic patterns. This algorithm provides a highly scalable methodology for comparative network biology, facilitating the systematic transfer of regulatory knowledge from fundamental model organisms to complex agricultural systems.

Despite its robust performance, the current alignment algorithm has notable limitations that present avenues for future research. First, the algorithm relies on static GRN representations aggregated over time-series data, which collapses the dynamic nature of stress responses into a single topological snapshot. Future iterations could incorporate dynamic or temporal network alignment layers to capture the phase-specific rewiring of these modules. Second, the quality of the alignment is inherently bounded by the accuracy of the initial ortholog mapping and the upstream ensemble GRN inference. Integrating multi-omics modalities, such as single-cell RNA-seq or spatial transcriptomics, alongside ATAC-seq for direct chromatin accessibility evidence, could further refine the edge weights and reduce the search space. While this study focuses on plant drought response, the relaxed topological alignment is a condition-and species-agnostic algorithm. It holds broad applicability for uncovering conserved disease modules or developmental pathways across diverse eukaryotic kingdoms.

## Acknowledgments

This material is based upon work supported by the National Science Foundation under Grant No. 2243691. We would like to express our sincere gratitude to Dr. Luis Herrera-Estrella and Dr. Liqing Zhang for providing valuable feedback and suggestions.

## Data availability statement

The primary transcriptomic data sets analyzed during this study are publicly available in the NCBI BioProject database under the accession numbers PRJNA527782 (*Sorghum bicolor*), PRJNA483231 (*Zea mays*), and PRJNA1155997 (*Arabidopsis thaliana*). The custom Python/R source code for the relaxed topological alignment algorithm, along with the statistical analysis scripts used to produce the results presented in this manuscript, are publicly available on GitHub at https://github.com/EveZhang19/VDT\_Cross\_Species\_Alignment.

## Notes

### Competing Interest Statement

The authors have declared no competing interest.

## References

1. Davidson EH. The regulatory genome: Gene regulatory networks in development and evolution. Elsevier; 2010.

2. Huynh-Thu VA, Sanguinetti G. Gene regulatory network inference: An introductory survey. Gene Regulatory Networks: Methods and Protocols. 2019:1–23.

3. Clark C, Kalita J. A comparison of algorithms for the pairwise alignment of biological networks. Bioinformatics. 2014;30(16):2351–9.

4. Ma L, Shao Z, Li L, Huang J, Wang S, Lin Q, et al. Heuristics and metaheuristics for biological network alignment: A review. Neurocomputing. 2022;491:426–41.

5. Zhao ZQ, Huang DS. A mended hybrid learning algorithm for radial basis function neural networks to improve generalization capability. Applied Mathematical Modelling. 2007;31(7):1271–81.

6. Chen J, Wang Z, Huang J. SAMNA: Accurate alignment of multiple biological networks based on simulated annealing. Journal of Integrative Bioinformatics. 2024;20(4):20230006.

7. Fuchs M, Riesen K. Fast approximate maximum common subgraph computation. Pattern Recognition Letters. 2025;190:66–72.

8. Ciriello G, Mina M, Guzzi PH, Cannataro M, Guerra C. AlignNemo: A local network alignment method to integrate homology and topology. PloS One. 2012;7(6):e38107.

9. Kazemi E, Hassani H, Grossglauser M, Pezeshgi Modarres H. PROPER: Global protein interaction network alignment through percolation matching. BMC Bioinformatics. 2016;17:1–16.

10. Mamano N, Hayes WB. SANA: Simulated annealing far outperforms many other search algorithms for biological network alignment. Bioinformatics. 2017;33(14):2156–64.

11. Altarawy D, Eid FE, Heath LS. PEAK: Integrating curated and noisy prior knowledge in gene regulatory network inference. Journal of Computational Biology. 2017;24(9):863–73.

12. Fitzek-Campbell E, Psaroudakis D, Weisshaar B, Junker A, Bräutigam A. A sublethal drought and rewatering time course reveals intricate patterning of responses in the annual Arabidopsis thaliana. bioRxiv. 2025:2025–07.

13. He C, Du Y, Fu J, Zeng E, Park S, White F, et al. Early drought-responsive genes are variable and relevant to drought tolerance. G3: Genes, Genomes, Genetics. 2020;10(5):1657–70.

14. Varoquaux N, Cole B, Gao C, Pierroz G, Baker CR, Patel D, et al. Transcriptomic analysis of field-droughted Sorghum from seedling to maturity reveals biotic and metabolic responses. Proceedings of the National Academy of Sciences. 2019;116(52):27124–32.

15. Bolser D, Staines DM, Pritchard E, Kersey P. Ensembl Plants: Integrating tools for visualizing, mining, and analyzing plant genomics data. In: Plant Bioinformatics: Methods and Protocols. Springer; 2016. p. 115–40.

16. Hufford MB, Seetharam AS, Woodhouse MR, Chougule KM, Ou S, Liu J, et al. De novo assembly, annotation, and comparative analysis of 26 diverse maize genomes. Science. 2021;373(6555):655–62.

17. Lamesch P, Berardini TZ, Li D, Swarbreck D, Wilks C, Sasidharan R, et al. The Arabidopsis Information Resource (TAIR): Improved gene annotation and new tools. Nucleic Acids Research. 2012;40(D1):D1202–10.

18. Wheeler D, Bhagwat M. BLAST QuickStart: Example-driven web-based BLAST tutorial. In: Comparative Genomics. Springer; 2007. p. 149–75.

19. Camacho C, Coulouris G, Avagyan V, Ma N, Papadopoulos J, Bealer K, et al. BLAST+: Architecture and applications. BMC Bioinformatics. 2009;10(1):421.

20. Emms DM, Kelly S. OrthoFinder: Phylogenetic orthology inference for comparative genomics. Genome biology. 2019;20(1):238.

21. Bolger AM, Lohse M, Usadel B. Trimmomatic: A flexible trimmer for Illumina sequence data. Bioinformatics. 2014;30(15):2114–20.

22. Bray NL, Pimentel H, Melsted P, Pachter L. Near-optimal probabilistic RNA-seq quantification. Nature Biotechnology. 2016;34(5):525–7.

23. Soneson C, Love MI, Robinson MD. Differential analyses for RNA-seq: Transcript-level estimates improve gene-level inferences. F1000Research. 2016;4:1521.

24. Tian F, Yang DC, Meng YQ, Jin J, Gao G. PlantRegMap: Charting functional regulatory maps in plants. Nucleic Acids Research. 2020;48(D1):D1104–13.

25. Love MI, Huber W, Anders S. Moderated estimation of fold change and dispersion for RNA-seq data with DESeq2. Genome Biology. 2014;15:1–21.

26. Robinson MD, McCarthy DJ, Smyth GK. edgeR: A Bioconductor package for differential expression analysis of digital gene expression data. Bioinformatics. 2010;26(1):139–40.

27. Law CW, Chen Y, Shi W, Smyth GK. voom: Precision weights unlock linear model analysis tools for RNA-seq read counts. Genome Biology. 2014;15:1–17.

28. Finkle JD, Wu JJ, Bagheri N. Windowed Granger causal inference strategy improves discovery of gene regulatory networks. Proceedings of the National Academy of Sciences. 2018;115(9):2252–7.

29. Huynh-Thu VA, Irrthum A, Wehenkel L, Geurts P. Inferring regulatory networks from expression data using tree-based methods. PLoS One. 2010;5(9):e12776.

30. Huynh-Thu VA, Geurts P. dynGENIE3: Dynamical GENIE3 for the inference of gene networks from time series expression data. Scientific Reports. 2018;8(1):3384.

31. Cirrone J, Brooks MD, Bonneau R, Coruzzi GM, Shasha DE. OutPredict: Multiple datasets can improve prediction of expression and inference of causality. Scientific Reports. 2020;10(1):6804.

32. Marbach D, Prill RJ, Schaffter T, Mattiussi C, Floreano D, Stolovitzky G. Revealing strengths and weaknesses of methods for gene network inference. Proceedings of the National Academy of Sciences. 2010;107(14):6286–91.

33. Marbach D, Schaffter T, Mattiussi C, Floreano D. Generating realistic in silico gene networks for performance assessment of reverse engineering methods. Journal of Computational Biology. 2009;16(2):229–39.

34. Prill RJ, Marbach D, Saez-Rodriguez J, Sorger PK, Alexopoulos LG, Xue X, et al. Towards a rigorous assessment of systems biology models: The DREAM3 challenges. PloS ONE. 2010;5(2):e9202.

35. Schaffter T, Marbach D, Floreano D. GeneNetWeaver: In silico benchmark generation and performance profiling of network inference methods. Bioinformatics. 2011;27(16):2263–70.

36. Zadrozny B, Elkan C. Obtaining calibrated probability estimates from decision trees and naive Bayesian classifiers. In: International Conference on Machine Learning. vol. 1; 2001.

37. Ding B, Liang M, Shi Y, Zhang R, Wang J, Huang Y, et al. The transcription factors DOF4. 6 and XND1 jointly regulate root hydraulics and drought responses in Arabidopsis. The Plant Cell. 2025;37(4):koaf083.

38. Ding B, Shi Y, Zhang R, Liang M, Sun X, Huang Y, et al. XND1-centered network regulates salt tolerance by integrating root xylem plasticity and Na+ unloading in Arabidopsis. Proceedings of the National Academy of Sciences. 2025;122(43):e2520667122.

